# The dark side of theoretical ecology

**DOI:** 10.1101/008813

**Authors:** Lev V. Kalmykov

**Affiliations:** Institute of Theoretical and Experimental Biophysics, Russian Academy of Sciences, Pushchino, Moscow Region, 142290 Russian Federation

## Abstract

Good science must be clearly transparent in its theories, models and experiments. Earlier David Tilman drew attention to the fact that ecologists investigate interspecific competition phenomenologically, rather than mechanistically. To create a mechanistic model of a complex dynamic system we need to logically describe interactions of its subsystems which lead to emergence of new properties on the macro-level. There are black-box, grey-box and white-box models of complex systems. Black-box models are completely nonmechanistic. We cannot investigate interactions of subsystems of such non-transparent model. A white-box model of a complex system has “transparent walls” and directly shows underlined mechanistic mechanisms – all events at micro-, meso-and macro-levels of the modeled dynamic system are directly visible at all stages. Grey-box models are intermediate. Basic ecological models are of black-box type, e.g. Malthusian, Verhulst, Lotka-Volterra models. These models are not individual-based and cannot show features of local interactions of individuals of competing species. That is why they principally cannot provide a mechanistic insight into interspecific competition. To create a white-box model we need a physical theory of the object domain and its intrinsic axiomatic system. On the basis of axiomatic system there is a possibility to logically generate a new knowledge by logical deterministic cellular automata. Understanding of biodiversity mechanisms is the global research priority. Only knowledge of mechanisms of interspecific interactions can allow us to efficiently operate in the field of biodiversity conservation. Obviously that such knowledge must be based on mechanistic models of species coexistence. In order to create a serviceable theory of biodiversity it is necessary to renew attempts to create a basic mechanistic model of species coexistence. But the question arises: *Why ecological modelers prefer to use the heaviest black-box mathematical methods which cannot produce mechanistic models of complex dynamic systems in principle, and why they do not use simple and long-known pure logical deterministic cellular automata, which easily can produce white-box models and directly generate clear mechanistic insights into dynamics of complex systems?*

---

Good science must be transparent in its theories, models and experiments. In my own research I often remember David Tilman’s great article which draws attention to the fact that ecologists investigate interspecific competition phenomenologically, rather than mechanistically (*2*). The article was published in 1987, however it is relevant for biodiversity science and mathematical modeling of complex systems even today. It discusses a problem among field experiments designed to test for the existence of interspecific competition in natural communities. Tilman suggests, ‘ *The design of the experiments, though, is a memorial to the extent to which the often-criticized Lotka-Volterra competition equations still pervade ecological thought. The experiments used a nonmechanistic, Lotka-Volterra-based, phenomenological definition of competition: two species compete when an increase in the density of one species leads to a decrease in the density of the other, and vice versa*. … *With a few notable exceptions, most ecologists have studied competition by asking if an increase in the density of one species leads to a decrease in the density of another, without asking how this might occur.* … *Experiments that concentrate on the phenomenon of interspecific interactions, but ignore the underlying mechanisms, are difficult to interpret and thus are of limited usefulness*.’(*2*) To design an adequate field experiment we should have a mechanistic model based on a mechanistic definition of interspecific competition. Otherwise, we will not be able to overcome the limitations of phenomenological approach which hides from us internal functional mechanisms of ecosystems. Only a mechanistic approach will allow us not only to constate the loss of biodiversity, but to understand what needs to be done to save it. How to create such a mechanistic model? First, we need to know how to mechanistically model a complex dynamic system. A complex dynamic system may be considered as consisting of subsystems that interact. Interactions between subsystems lead to the emergence of new properties, e.g. of a new pattern formation. Therefore we should define these subsystems and logically describe their interactions in order to create and investigate a mechanistic model. If we want to understand how a complex dynamic system works, we must understand cause-effect relations and part-whole relations in this system. The causes should be sufficient to understand their effects and the parts should be sufficient to understand the whole. There are three types of possible models for complex dynamic systems: black-, grey-, and white-box models (Fig. 1).

**Figure 1.**
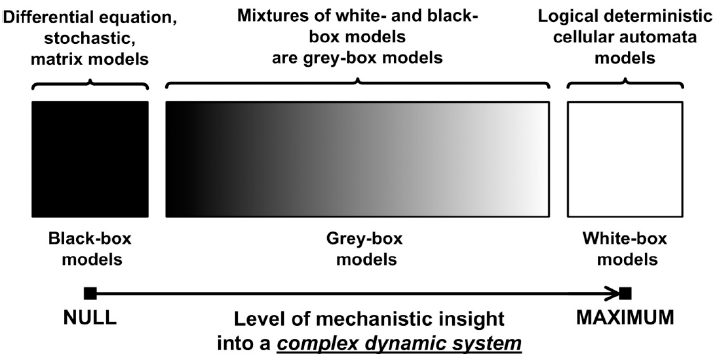
Three types of possible models for complex dynamic systems.

Black-box models are completely nonmechanistic. We cannot investigate interactions of subsystems of such a non-transparent model. A white-box model of complex dynamic systems has ‘transparent walls’ and directly shows underlying mechanisms – all events at micro-, meso-and macro-levels of a modeled dynamic system are directly visible at all stages. Logical deterministic cellular automata is the only known approach, which allows to create white-box models of complex dynamic systems (*3*). A micro-level is modeled by a lattice site (cellular automata cell). A meso-level of local interactions of micro-objects is modeled by a cellular automata neighbourhood. A macro-level is modeled by the entire cellular automata lattice. Unfortunately, this simple approach is commonly used in the overloaded form, what makes it less transparent. This is achieved by adding differential equations and stochasticity. Grey-box models are intermediate and combine black-box and white-box approaches. Basic ecological models are of black-box type, e.g. Malthusian, Verhulst, Lotka-Volterra models. These models are not individual-based and cannot show features of local interactions of individuals of competing species. That is why they principally cannot provide a mechanistic insight into interspecific competition.

A white-box model of a complex system is completely mechanistic. A white-box modelling is axiomatic modelling. To begin to create a white-box model we need to formulate an intrinsic axiomatic system based on a general physical understanding of the subject area under study. Axioms are first principles of the subject. René Descartes proposed that axiomatic inference is universal for any science on condition that a system of axioms is complete and provided that axioms are unquestionably true, clear and distinct (*4*). Descartes was inspired by Euclidean geometry which investigates the relations between ideal spatial figures. When scientists verify a theory first of all they should strictly verify its axioms. If at least one axiom is inadequate or an axiomatic system is incomplete, then the theory is inadequate too (*5*). Let’s consider an example of the inadequacy of ecological models in result of incompleteness of their axiomatic system. There are many models of population dynamics that do not take into account what happens with individuals after their death. Dead individuals instantly disappear with roots, stubs, etc. ‘*One reason for the lack of understanding on the part of most botanists results from their failure to take into account the phenomenon of regeneration in plant communities, which was first discussed in general terms by A. S. Watt in 1947*.’ (*6*)

Stephen Hubbell in his Unified Neutral Theory of Biodiversity (UNTB) in fact refuses a mechanistic understanding of interspecific competition: ‘*We no longer need better theories of species coexistence; we need better theories for species presence-absence, relative abundance and persistence times in communities that can be confronted with real data. In short, it is long past time for us to get over our myopic preoccupation with coexistence*’ (*7*). However, he admits that *‘the real world is not neutral’* (*8*). Since the basic postulate (axiom) of the UNTB about ecological neutrality of similar species is wrong, this theory cannot be true. In addition, local interactions of individuals are absent in the neutral models in principle. That is why neutral models cannot provide a mechanistic insight into biodiversity. The UNTB models are of black-box and dark grey-box types only – Fig.1. I agree with James Clark, that the dramatic shift in ecological research to focus on neutrality distracts environmentalists from the study of real biodiversity mechanisms and threats (*9*). Within the last decade, the neutral theory has become a dominant part of biodiversity science, emerging as one of the concepts most often tested with field data and evaluated with models (*9*). Neutralists are focused on considering unclear points of the neutral theory – the ecological drift, the link between pattern and process, relations of simplicity and complexity in modelling, the role of stochasticity and others, but not the real biodiversity problems themselves (*8*). Attempts to understand neutrality instead of biodiversity understanding look like attempts to explain the obscure by the more obscure. Nonmechanistic models make it difficult to answer basic ecological questions, e.g. Why are there so many closely allied species? (*10*) An example of the difficult ecological discussion is the debates ‘Ecological neutral theory: useful model or statement of ignorance?’ on the forum Cell Press Discussions (*11*).

Understanding of mechanisms of interspecific coexistence is a global research priority. These mechanisms can allow us to efficiently operate in the field of biodiversity conservation. Obviously, such knowledge must be based on mechanistic models of species coexistence. In order to create a practically useful theory of biodiversity, it is necessary to renew attempts to create a basic mechanistic model of species coexistence. But the question arises: *Why do ecological modelers prefer to use the heaviest black-box mathematical methods, which cannot produce mechanistic models of complex dynamic systems in principle, and not use simple and long-known pure logical deterministic cellular automata, which easily can produce white-box models and directly obtain clear mechanistic insights into dynamics of complex systems?*

## Acknowledgements

I thank Vyacheslav L. Kalmykov for useful discussions and suggestions.

I thank Kylla M. Benes for helpful suggestions and edits.

## References

1. J. Maddox, The dark side of molecular biology. Nature 363, 13 (1993). doi: http://dx.doi.org/10.1038/363013a0

2. D. Tilman, The importance of the mechanisms of interspecific competition. The American Naturalist 129, 769 (1987). doi: http://dx.doi.org/10.1086/284672

3. L. V. Kalmykov, V. L. Kalmykov, Verification and reformulation of the competitive exclusion principle. Chaos, Solitons & Fractals 56, 124 (2013). doi: http://dx.doi.org/10.1016/j.chaos.2013.07.006

4. R. Descartes, Discourse on the method of rightly conducting one’s reason and of seeking truth in the sciences. D. A. Cress, Ed., (Hackett Pub. Co., Indianapolis, 1637/1980), 42 p.

5. B. Spinoza, Principles of Cartesian Philosophy: And, Metaphysical Thoughts. (Hackett Pub. Co., 1998).

6. P. J. Grubb, The maintenance of species-richness in plant communities: the importance of the regeneration niche. Biological Reviews 52, 107 (1977). doi: http://dx.doi.org/10.1111/j.1469-185X.1977.tb01347.x

7. S. P. Hubbell, The unified neutral theory of biodiversity and biogeography. Monographs in population biology; 32 (Princeton University Press, Princeton, N.J. ; Oxford, 2001), pp. xiv, 375 p.

8. J. Rosindell, S. P. Hubbell, F. He, L. J. Harmon, R. S. Etienne, The case for ecological neutral theory. Trends in Ecology & Evolution 27, 203 (2012). doi: http://dx.doi.org/10.1016/j.tree.2012.01.004

9. J. S. Clark, Beyond neutral science. Trends in Ecology & Evolution 24, 8 (Jan, 2009). doi: http://dx.doi.org/10.1016/j.tree.2008.09.004

10. Anonymous. British Ecological Society: Easter Meeting 1944: Symposium on “The Ecology of Closely Allied Species”. Journal of Animal Ecology 13, 176 (1944). Stable URL: http://www.jstor.org/stable/1450

11. P. Craze, Ecological neutral theory: useful model or statement of ignorance? Available from Cell Press Discussions: < http://news.cell.com/discussions/trends-in-ecology-and-evolution/ecological-neutral-theory-useful-model-or-statement-of-ignorance > (2012).

